# Iron deficiency in people with obesity drives defective Natural Killer cell mitochondrial fitness and function

**DOI:** 10.1101/2024.01.10.575005

**Authors:** Conor De Barra, Eimear Ryan, Michelle Sugrue, Odhran Ryan, Evelyn Lynn, Helen M. Heneghan, Cormac McCarthy, Paul N. Moynagh, Linda V. Sinclair, Nicholas Jones, Andrew E. Hogan, Donal O’Shea

**Author notes:** Corresponding Author **Address for correspondence:** Dr Andrew Hogan –, Address: Biosciences Building, Maynooth University, Maynooth, Co. Kildare, Ireland. Joint senior authorship (Andrew E. Hogan, Donal O’Shea).

## Abstract

Natural killer (NK) cells are a population of innate effector lymphocytes, involved in host-defences against viral infections and cancer. Upon activation, NK cells can produce a milieu of cytotoxic molecules and cytokines, which can directly target infected and transformed cells, but also amplify an immune response. Metabolic rewiring underpins NK cell effector functionality, providing the required signals, energy and biointermediates to support their immune responses. Obesity is associated with significant defects in the functionality of human NK cells, especially in the periphery. Dysregulated cellular metabolism has been demonstrated to be a major mechanistic driver of the reported defects. However, how obesity links to defective NK cell metabolism and functionality remains unclear. Iron deficiency is a common co-morbidity in people living with obesity (PWO). Recent studies have highlighted the importance for iron in host immunity, with murine models of iron deficiency resulting in defective cellular metabolism and function. We hypothesized that obesity-driven iron deficiency might underpin the reported defects in NK cells. Our data demonstrates that in response to cytokine stimulation, healthy human NK cells utilize iron to support their metabolic activity and cytokine responses. In a cohort of PWO, we demonstrate alterations in NK cell metabolism, mitochondrial fitness and cytokine production. Furthermore, upon stratification into PWO with normal iron status versus low iron status, we show the observed obesity-related defects in NK cell metabolism, mitochondrial fitness and cytokine production are concentrated in the PWO with low-iron status. Collectively, our data highlights the importance of iron for human NK cell responses and provides evidence that obesity-driven defects in NK cell metabolism and function are linked in part to altered iron availability.

## Introduction

Natural killer (NK) cells are a population of innate effector lymphocytes involved in immunity against viral infections and cancers^1, 2^. NK cells are armed with cytotoxic molecules capable of lysing transformed or infected target cells^3, 4, 5^. Upon cytokine stimulation NK cells can also produce a multitude of inflammatory cytokines and chemokines^6^, and thus are capable of initiating and/or amplifying a stronger immune response. Underpinning this robust NK cell effector profile is an immunometabolic program which provides them with the signals, energy and biosynthetic intermediates to produce effector molecules such as interferon gamma (IFN-γ)^7^. At rest, peripheral NK cells are quiescent cells with low metabolic rates^8, 9^, whereas cytokine stimulated NK cells display elevated rates of glycolysis and oxidative phosphorylation (OxPHOS)^10,11^. This increased metabolic activity is paired with an increased demand for nutrients including glucose and amino acids^8, 11, 12, 13^. Previous work from our group has highlighted defects in NK cell metabolism ^14, 15, 16, 17^ as a mechanism underpinning reported defects in NK cell functionality in people with obesity (PWO)^18, 19, 20, 21^. Altered systemic lipid profiles was proposed as an obesogenic driver of the defects in NK cells cellular metabolism^21^. However, obesity is associated with significant alterations in systemic nutrient availability, with one of the most well defined being iron^22^. Recently, several studies have highlighted the importance of iron, an essential trace element, for host immune responses and homeostasis ^23, 24, 25^. Iron is essential for several critical cellular processes such as OxPhos, DNA synthesis and cell proliferation ^26, 27, 28^. Currently, it is unknown if lower iron availability in people with obesity is linked to the reported defects in NK cells. In the current study, we demonstrate that in healthy human NK cells, after cytokine stimulation, extracellular iron supports protein translation, adenosine triphosphate (ATP) production, and cytokine responses. In the setting of obesity, we show a loss of mitochondrial fitness and reduced cytokine responses is underpinned by altered iron availability. Collectively, our data demonstrates that human NK cells require extracellular iron for their metabolic and functional responses after cytokine stimulation, and altered iron availability in obesity is associated with defective NK cells.

## RESULTS

### Cytokine activated NK cells increase expression of CD71 and transferrin uptake

First, we investigated if human NK cells increase their expression of the transferrin receptor CD71 upon cytokine stimulation (IL-12/IL-15) and observed a significant increase in both the percentage of NK cells expressing CD71, and the mean fluorescent intensity (MFI) of CD71 expression (Figure 1A-D). Next, we investigated if cytokine activated NK cells increase their uptake of transferrin, using a transferrin uptake assay, and demonstrate increases in both the percentage and MFI of transferrin uptake (Figure 1E-G). To extend our investigations into the setting of obesity, we recruited a cohort of people with obesity (PWO) and confirmed altered iron availability, with lower serum iron and transferrin saturation in comparison to healthy controls (Figure 1H-I). Next, we investigated CD71 expression and transferrin uptake in this cohort of PWO. There was tendency for NK cells from PWO to show higher expression of CD71 and while this did not reach statistical significance, these cells demonstrated elevated basal transferrin uptake in NK cells when compared to healthy controls (Figure 1J-K).

**Figure 1:**
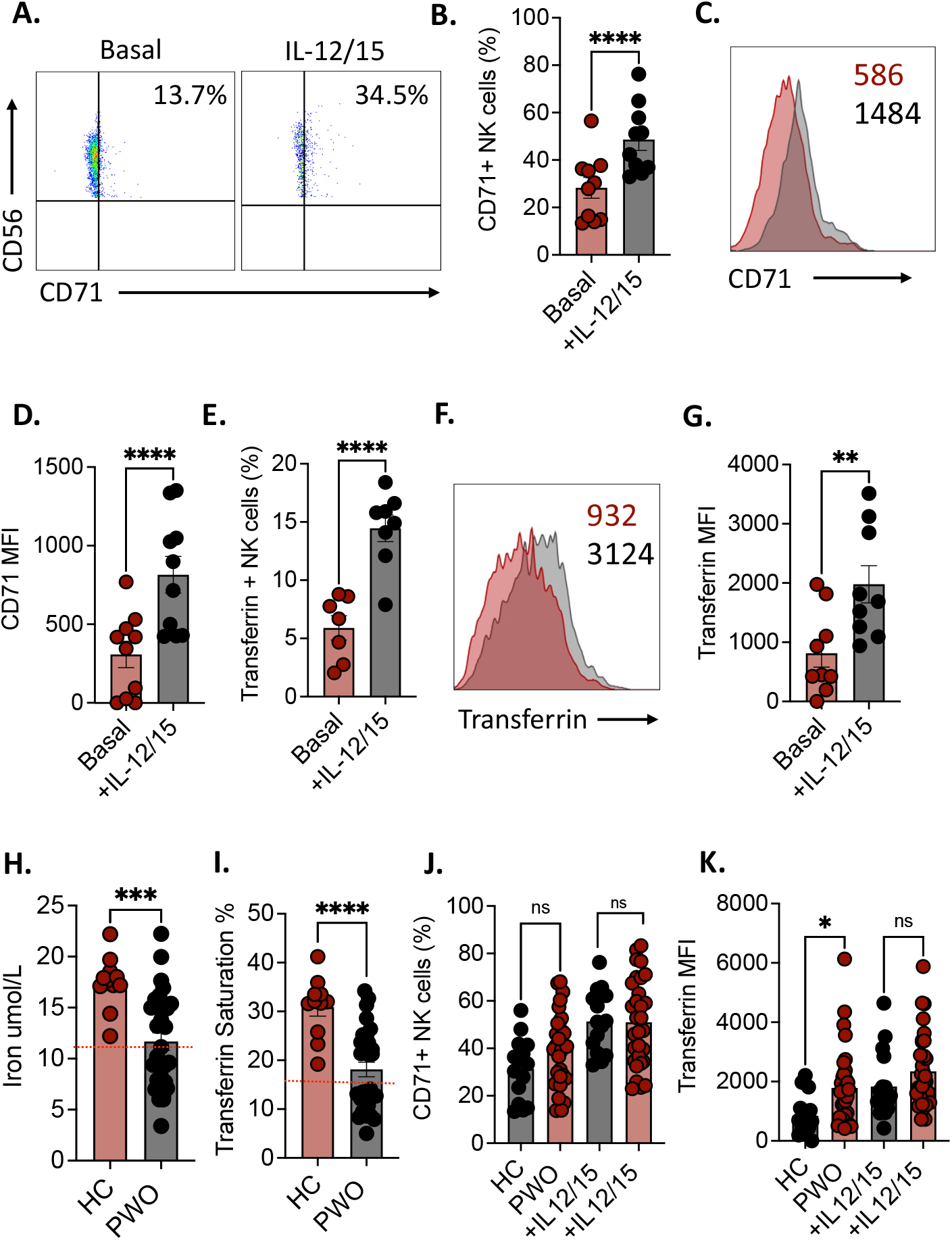
Cytokine activated NK cells increase expression of CD71 and transferrin uptake. (A) Representative dot plot showing frequencies of CD71 expressing NK cells. (B) Bar chart detailing the frequencies of CD71 expressing NK cells, either basal or stimulated with IL-12/15 for 18 hours. (C) Representative histogram showing MFI of CD71 on NK cells either basal or stimulated with IL-12/15 for 18 hours. (D) Bar chart showing the frequencies of CD71 expressing NK cells either basal or stimulated with IL-12/15 for 18 hours. (E-G) Bar chart and representative histogram detailing the frequency of transferring uptake and MFI of transferrin in NK cells either basal or stimulated with IL-12/15 for 18 hours. (H-I) Bar charts showing the iron content (umol/L) and Transferrin Saturation (%) of blood from cohorts of healthy controls (HC) and people with obesity (PWO). (J) Bar chart showing the frequencies of CD71 expressing NK cells from both HC and PWO, either basal or stimulated with IL-12/15 for 18 hours. (K) Bar chart displaying the frequencies of transferrin uptake in NK cells from both HC and PWO, either basal or stimulated with IL-12/15 for 18 hours. ns, no significance; *p < 0.05; **p < 0.01; ***p < 0.001.

### Cytokine activated human NK cells require iron for optimal metabolic responses

We next determined whether human NK cells require extracellular iron to support metabolic activity after cytokine stimulation. We utilized SCENITH^29^, a single cell approach which allows immunometabolic characterisation based on cellular protein translation. We show that NK cells increased their rates of protein translation, as measured by puromycin incorporation after cytokine stimulation (Figure 2A-B). Furthermore, upon stimulation NK cells decreased their reliance on OxPhos relying more on glycolysis (Figure 2C-D). Next, using the iron chelator deferoxamine (DFO) to model reduced iron availability, we show that NK cell protein translation is reduced under low iron conditions (Figure 2E-F). We also demonstrate that NK cell metabolic dependencies change under low iron conditions, with increased glucose dependency and reduced OxPhos dependency (Figure 2G-H). Mirroring the effects of DFO on NK cells, our cohort of PWO displayed reduced capacity for protein translation with significantly reduced puromycin incorporation (Figure 2I-J). While there is a significant reduction in puromycin incorporation, NK cells from PWO demonstrate similar glycolytic and OxPhos dependencies to NK cells from healthy controls (Figure 2 K-L).

**Figure 2:**
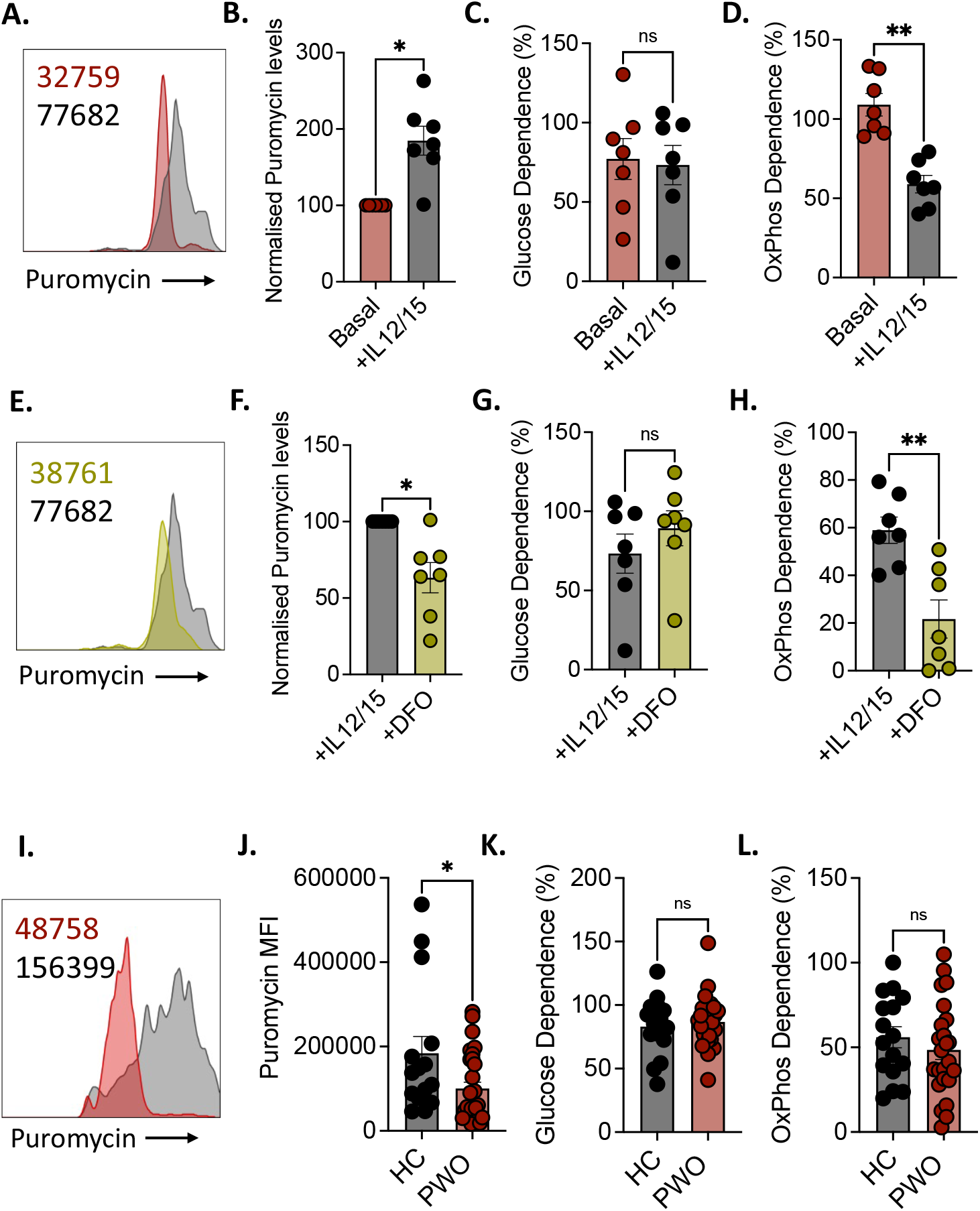
Cytokine activated NK cells require iron for metabolic and functional responses. (A) Representative histogram detailing MFI of puromycin incorporation into NK cells either basally or stimulated with IL-12/15 for 18 hours. (B) Bar chart showing change in puromycin MFI relative to basal of NK cells stimulated with IL-12/15 for 18 hours. (C-D) Bar charts showing the metabolic dependence of NK cells on glycolysis and OxPhos respectively either basal or stimulated with IL-12/15 for 18 hours. (E) Representative histogram detailing MFI of puromycin of NK cells stimulated with IL-12/15 for 18 hours, in the absence or presence of the iron chelator DFO. (F) Bar chart showing change in puromycin MFI relative to stimulated NK cells or NK cells in the presence of DFO for 18 hours. (G-H) Bar charts showing the metabolic dependence of NK cells on glycolysis and OxPhos respectively, stimulated with IL-12/15 for 18 hours in the presence or absence of DFO. (I) Representative histogram detailing MFI of puromycin of NK cells, stimulated with IL-12/15 for 18 hours either from healthy controls (HC) or people with obesity (PWO). (J) Bar chart showing puromycin MFI of NK cells stimulated with IL-12/15 for 18 hours, from either PWO or HC. (K-L) Bar charts showing the metabolic dependence of NK cells on glycolysis and OxPhos respectively, stimulated with IL-12/15 for 18 hours, either from PWO or HC. ns, no significance; *p < 0.05; **p < 0.01; ***p < 0.001.

### Altered iron availability drives reduced mitochondrial fitness in human NK cells

With the noted reduction in NK cell mitochondrial metabolism under conditions of limited iron, we next investigated the impact of reduced iron availability on human NK cell mitochondrial mass and polarisation. We demonstrate an accumulation of dysregulated mitochondria, as measured by an increase in the proportion of NK cells with high MitoTracker Green and low MitoTracker Deep Red, indicating large but depolarised mitochondria^30^, after DFO treatment (Figure 3A-B). This was reinforced with a decrease in the ratio of MitoTracker Green to MitoTracker Deep Red in DFO treated NK cells (Figure 3C). We next assessed the impact of reduced iron availability on NK cell ATP production and found a marked reduction in ATP production by NK cells treated with DFO (Figure 3D). This was also associated this was paired with diminished ability to produce IFNγ after cytokine stimulation (Figure 3E). Mirroring DFO treated NK cells, peripheral NK cells from PWO displayed similar phenotype with increased mitochondrial dysfunction leading to significantly reduced ATP and IFNγ production (Figure 3F-I).

**Figure 3:**
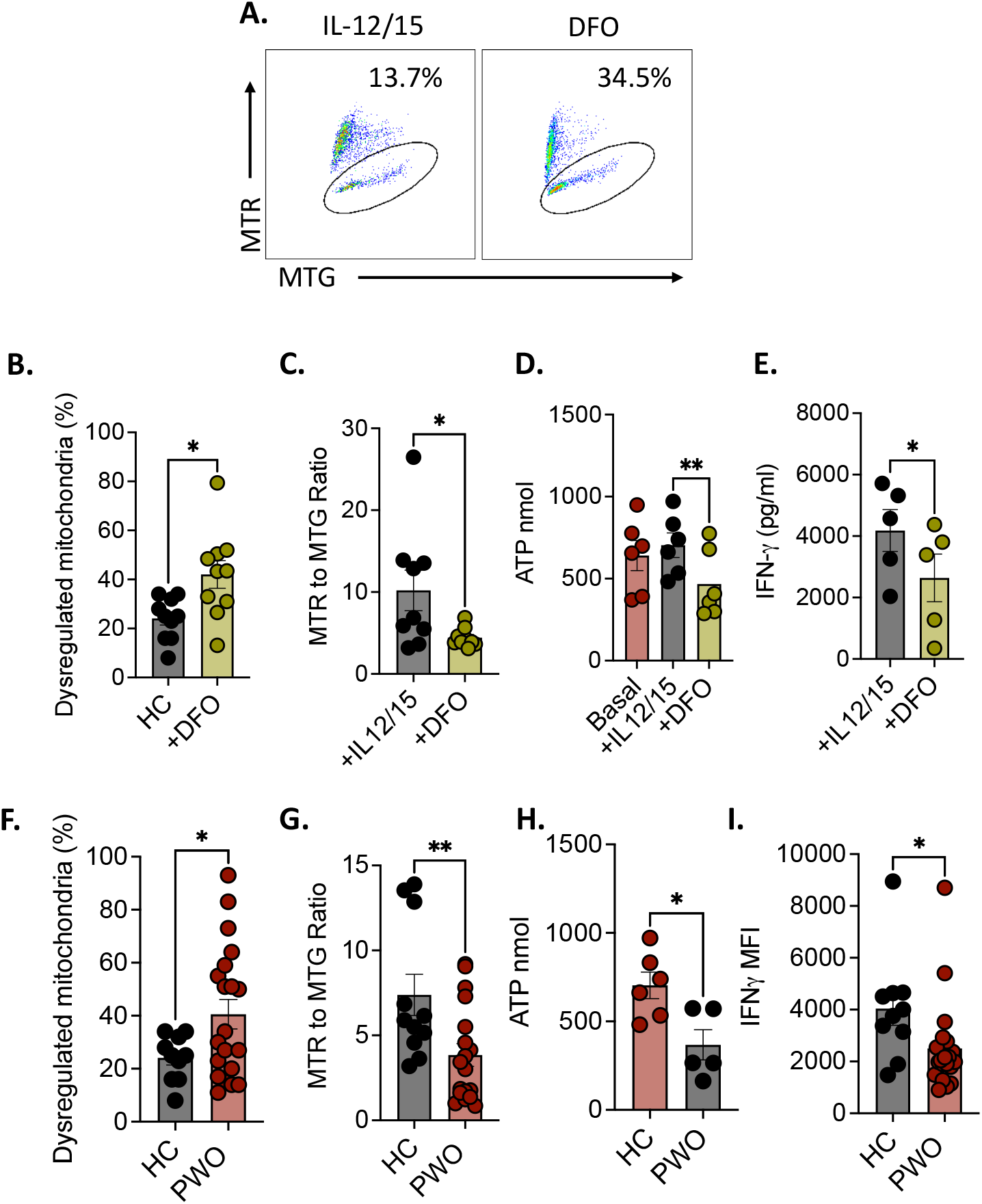
Mitochondrial function is dependent on iron availability. (A) Representative dot plots showing frequencies of dysregulated mitochondria (high Mitotracker green & low Mitotracker DeepRed) in NK cells, stimulated with IL-12/15 for 18 hours, in the presence or absence of DFO. (B) Bar chart showing frequencies of dysregulated mitochondria in NK cells, stimulated with IL-12/15 for 18 hours, in the presence or absence of DFO. (C) Bar chart describing the ratio of MitoTracker Deep Red MFI to MitoTracker Green MFI of NK cells, stimulated with IL-12/15 for 18 hours, in the presence or absence of DFO. (D) Bar chart detailing the production of ATP (nmol) in NK cells, either basally, or stimulated with IL-12/15 for 18 hours, in the presence or absence of DFO. (E) Bar chart detailing the levels of IFNγ produced by NK cells stimulated with IL-12/15 for 18 hours, in the presence or absence of DFO. (F) Bar chart describing frequencies of dysregulated mitochondria in NK cells, from HC or PWO, stimulated with IL-12/15 for 18 hours. (G) Bar chart showing the ratio of MitoTracker Deep Red MFI to MitoTracker Green MFI of NK cells, from HC or PWO, stimulated with IL-12/15 for 18 hours. (H) Bar chart detailing the production of ATP (nmol) in NK cells, from HC or PWO, stimulated with IL-12/15 for 18 hours. (I) Bar chart detailing IFNγ MFI in NK cells stimulated with IL-12/15 for 18 hours, from either HC or PWO. ns, no significance; PWO, people with obesity. *p < 0.05; **p < 0.01; ***p < 0.001.

### Low iron status in obesity is associated with defective NK cell metabolism and functionality

With the parallels between healthy NK cells treated with DFO and NK cells from PWO, we next aimed to established if the dysregulations observed were directly linked to altered iron availability, by stratifying our PWO cohort into those with low (<15%) transferrin saturation (Low TS) and those in the normal (15% to 50%) transferrin saturation range (Normal TS)^31, 32^ (Figure 4A). Interestingly, PWO with low TS expressed significantly more CD71 than those with normal TS (Figure 4B). Next, we investigated metabolic activity, and show that cytokine stimulated NK cells from the low TS cohort had reduced rates of protein translation and mitochondrial fitness in comparison to those in the normal TS cohort (Figure 4C-E). To investigate if this loss of mitochondrial fitness underpinned the defective functional of NK cells from PWO, we performed statistical analysis, and show a strong correlation between the proportion of dysregulated mitochondria and the percentage of IFNγ producing cells (Figure 4F). We also demonstrate that transferrin saturation levels in PWO are positively linked to the IFNγ producing ability of cytokine stimulated NK cells (Figure 4G). Supporting this observation, we also found that the IFNγ levels in NK cells from low TS cohort were significantly lower than healthy controls, whereas no significance was found between normal TS cohort and controls (Figure 4H).

**Figure 4:**
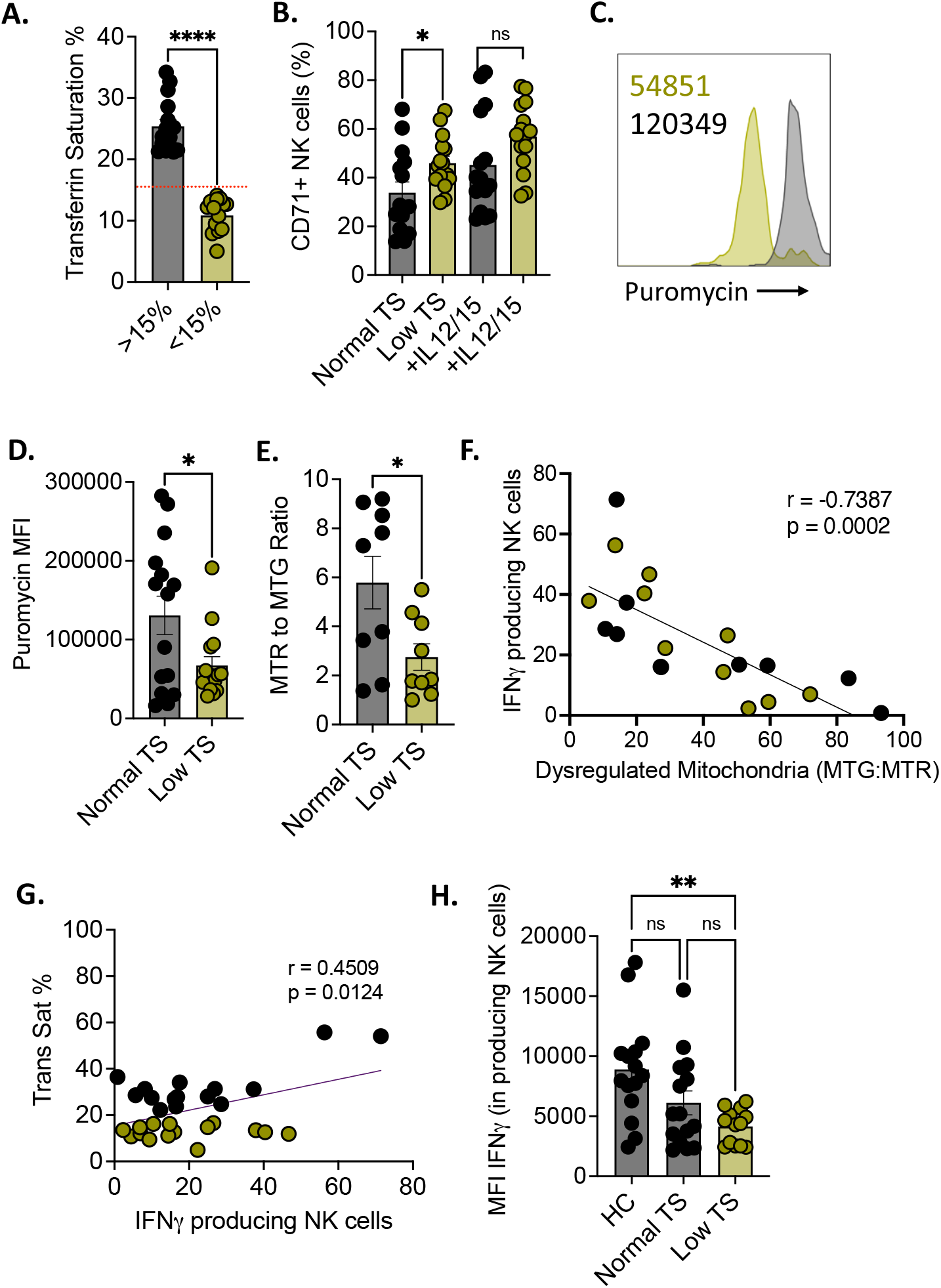
Low iron status in obesity is associated with defective NK cell metabolism and functionality. (A) Bar chart detailing the transferrin saturation (%) of PWO, dividing the cohort into those with normal transferrin saturation and low transferrin saturation. (B) Bar chart showing the frequencies of CD71 expressing NK cells from PWO, both normal TS and low TS, either basal or stimulated with IL-12/15 for 18 hours. (C) Representative histogram detailing MFI of puromycin of NK cells stimulated with IL-12/15 for 18 hours, from PWO with either normal TS or low TS. (D) Bar chart showing puromycin MFI of NK cells, stimulated with IL-12/15 for 18 hours, from PWO with either normal TS or low TS. (E) Bar chart detailing the ratio of MitoTracker Deep Red MFI to MitoTracker Green MFI of NK cells, from PWO, both normal TS and low TS, either basal or stimulated with IL-12/15 for 18 hours. (F) Scatter plot describing the relationship between the percentage of IFN-γ producing NK cells and the percentage of dysregulated mitochondria in both the normal TS group and low TS group. (G) Scatter plot detailing the relationship between the percentage of IFN-γ producing NK cells and the transferrin saturation in both the normal TS group and low TS group. (H) Bar chart showing the MFI of IFN-γ from IFN-γ producing NK cells from HC, normal TS and low TS cohorts stimulated with IL-12/15 for 18 hours. ns, no significance, *p < 0.05; **p < 0.01; ***p < 0.001.

## Discussion

Obesity is associated with significant changes in circulating NK cell metabolism and functionality ^14, 15, 17, 33, 34^. However, the specific factors that underpin these changes remain unclear. One of the most common systemic nutritional alterations in obesity is reduced iron availability^22^, yet the role of iron and the effect of iron deficiency on human NK cells remain unclear. In this study, we demonstrate that exogenous iron is critical for human NK cell metabolic fitness and IFNγ production. We provide evidence which links low iron availability to dysfunctional NK cells in PWO.

Iron is critical for numerous basic cellular processes including bioenergetics, with many proteins in these processes requiring iron to function^35^. There are various forms of iron found in the body with varying degrees of bioavailability, with the vast majority of iron stored in tissues, especially the liver, as ferritin^36^. In the bloodstream iron is bound to transferrin, and this complex is transported to various organs and tissues^35^. Transferrin bound iron is the transported into target cells via the transferrin receptor CD71^37^. We investigated the expression of CD71 on human NK cells, and while basal expression was low, cytokine stimulation of these cells caused strong upregulation of CD71. This supported previous studies in murine NK cells, which also noted robust increases in CD71 expression after cytokine stimulation ^24^. Interestingly the upregulation of CD71 on activated NK cells mirrors equivalent CD71 upregulation in functionally related CD8^+^ T cells upon activation^38, 39^.

In our cytokine stimulated human NK cells, the increase in CD71 expression was also associated with an increase in transferrin uptake, again supporting studies in mice which demonstrated transferrin uptake by NK cells^24, 40^. In our cohort of PWO, we report reduced serum iron and transferrin saturation, in line with previous meta-analysis^22^. In this cohort we noted no significant overall difference in CD71 expression by peripheral NK cells, however when we further stratified our obesity cohort into those with normal iron availability and those with reduced iron availability, we noted a marked increase in basal CD71 expression in the reduced iron cohort. This potential compensatory mechanism of elevated CD71 was also observed in ILC3 under iron limited conditions^25^.

As outlined, iron is critical for cellular metabolism, in particular mitochondrial metabolism^41^. We found that limiting iron availability using the iron chelator deferoxamine (DFO) reduced the rates of protein translation in cytokine activated NK cells, that likely is due to decrease in metabolic activity given that over half of the energy produced by cell metabolism is utilised in protein biosynthesis. Using SCENITH ^29^, we observed a significant decrease in OxPhos dependent protein translation, recapitulating reduced OxPhos rates in ILC3 treated with DFO^25^. In line, with these observations, NK cells from PWO displayed reduced levels of protein synthesis, with the greatest reductions in the individuals with low iron availability, potentially linking low iron status to defective NK cell metabolism in people with obesity. In CD8+ T cells, iron deficiency resulted in dysregulated mitochondria and reduced ATP production^23^, so we investigated this in healthy human NK cells, and found that DFO treatment resulted in an accumulation of dysregulated depolarized mitochondria^30^, which was paired with a significant drop in ATP production. Strikingly, the DFO driven phenotype of poor mitochondrial fitness (depolarized mitochondria and reduced ATP production) phenocopied NK cells from PWO, reinforcing the concept that obesity-related defects in NK cell metabolism may be linked to altered iron availability.

Metabolism is intrinsically linked to NK cell functionality, including the ability to produce key cytokines such as IFNγ^9^, which is required for NK cell mediated anti-viral responses^42, 43^, and can directly inhibit viral replication and drive malignant cell apoptosis ^44, 45, 46^. NK cell mitochondrial fitness has been directly linked to their effector function including their IFNγ production^47, 48^. In line with these observations, we observed a very strong correlation between mitochondrial fitness and IFNγ production in people with obesity. Similarly, we found that the PWO with reduced iron availability had greater reductions in the frequencies of IFNγ producing NK cells, again highlighting the importance of iron for NK cell fitness. This concept is further strengthened by data from Santosa and colleagues, who report defective adaptive NK cell responses in a murine model of iron deficiency^40^.

In summary, our work demonstrates that human NK cells require iron for their metabolic fitness and effector function. We show that in the setting of obesity, low iron availability is strongly linked to defective NK cell metabolism and cytokine production. This study provides new insight into factors underpinning obesity-driven defects in immunity, and presents a possible therapeutic target for improving NK cell responses in a group at greater risk of adverse outcomes with viral infection and cancer.

## Materials & methods

### Study cohorts & ethical approval

In total a cohort of 30 people living with obesity (BMI>30) were recruited – 15 individuals with normal transferrin saturation and 15 with low transferrin saturation. Both groups were matched for sex, age and BMI (see Table S1). After informed consent, a peripheral blood sample was taken for research purposes. Inclusion criteria included ability to give informed consent, 18-65 years of age, no current or recent (<2 weeks) infection, and no use of anti-inflammatory medications including GLP-1 analogue therapies. A cohort of 15 healthy adult donor were recruited as a control group. Ethical approval was obtained from both St Vincent’s University Medical Ethics Committee and Maynooth University Ethics Committee

### Preparation of peripheral blood mononuclear cells (PBMC) and flow cytometric analysis

PBMC samples were isolated by density centrifugation in SeptMate 50mL tube over Lymphoprep, from fresh peripheral blood samples. After using a viability stain, eBiosciences 506 Live Dead Stain (Invivogen), NK cells were identified as CD3^-^, CD56^+^ cells (Miltenyi BioTec). When appropriate NK cells were fixed and permeabilised using the True-Nuclear Transcription Factor Buffer set (BioLegend) according to the manufacturer’s instructions. Transferrin receptor levels were measured via CD71 (Miltenyi Biotec). Cell populations were acquired using a Attune NXT flow cytometer and analysed using FlowJo software (Treestar). Results are expressed as a percentage of the parent population as indicated and determined using flow minus-1 (FMO) and unstained controls.

### NK Cell Isolation

NK cells were isolated from PBMCs using a NK cell negative selection kit from Miltenyi Biotec as per manufacturer’s instructions using LS columns (Miltenyi Biotec) and MACS Buffer (Miltenyi Biotec). NK cell purity was confirmed via flow cytometry (>93% purity). Additionally, monocyte contamination was confirmed to be below 1%.

### NK cell SCENITH assay

PBMCs (2 × 10^6^ /mL) were activated using IL-15 (50ng/mL) and IL-12 (30ng/mL) for 18 hours. Cells were seeded into a 96-well plate, and treated as a control, or with 2-Deoxy-D-Glucose (100mM), Oligomycin (1μM), or both. Following incubation at 37°C for 15 minutes, cells were treated with Puromycin (11μM) and incubated for a further 25 minutes. Cells were washed with ice-cold PBS to stop puromycin incorporation. Cells were then stained for viability. NK cells were stained for extracellular markers and fixed, as outlined above. Staining of puromycin was achieved using anti-puromycin monoclonal antibody (AlexaFluor488, Sigma), in permeabilization buffer (BioLegend).

### NK cell transferrin uptake assay

Stimulated and unstimulated PBMCs (1 × 10^6^), activated using IL-15 (50ng/mL) and IL-12 (30ng/mL) for 18 hours, were rested in serum-free HPLM with 5% BSA for 2 hours. Cells were then washed in serum-free HPLM with 0.5% BSA and incubated with 5μg/ml Transferrin-AlexaFluor647 (Invitrogen) for 10 minutes at 37°C. Holo-transferrin (500μg/ml, Sigma-Aldrich) was used to competitively control for transferrin-uptake. An additional sample was stained at 4°C. Cells were washed in ice-cold HPLM with 0.5% BSA to stop membrane trafficking. Cells were then stained for viability, and NK cells were labelled for extracellular markers, to be analysed by flow cytometry.

### NK cell mitochondrial analysis

PBMCs (1 × 10^6^ /mL) were activated using IL-15 (50ng/mL) and IL-12 (30ng/mL) for 18 hours, in the absence or presence of DFO (200μM, Sigma). Cells were seeded into a 96-well plate and washed in serum-free buffer. Cells were then stained for viability as outlined above, and NK cells were stained for extracellular markers. Cells were then washed and stained with Mitotracker Deep Red FM (50μM, ThermoFisher Scientific) and Mitotracker Green (50μM, ThermoFisher Scientific) in PBS, and incubated for 1 hour at 37°C. Cells were subsequently analysed by flow-cytometric analysis.

### NK cell functional analysis

PBMCs (1 × 10^6^ /mL) were activated using IL-15 (50ng/mL) and IL-12 (30ng/mL) for 18 hours, in the absence or presence of DFO (200μM, Sigma). After using a viability stain, eBiosciences 506 Live Dead Stain (Invivogen), NK cells were identified as CD3^-^, CD56^+^ cells (Miltenyi BioTec). Cells were also stained for NKG2D (Miltenyi Biotec). The PBMCs were then fixed and permeabilised using the True-Nuclear Transcription Factor Buffer set (BioLegend) according to the manufacturer’s instructions. In perm buffer the cells were then stained for Granzyme B, IFN-γ and Perforin (Miltenyi Biotec).

### NK cell ATP assay

Isolated NK cells (1 × 10^6^ /ml) were activated using IL-15 (50ng/mL) and IL-12 (30ng/ml) for 18 hours. A luminescence ATP assay kit (abcam) was used. Reagents in the kit we reconstituted as per manufacturer’s instructions. A standard curve was also prepared as per the kit’s the instructions. NK cells were harvested and washed with PBS. 100μL of resuspended NK cells was added to a black walled, clear bottomed plate. 50μL of detergent was added to each well and the plate was placed on an orbital shaker for 5 minutes at 600-700 rpm. 50μ of substrate solution was added and the plate was returned to the orbital shaker for 5 minutes at 600-700 rpm. The plate was then covered and placed in the dark for 10 minutes before luminescence was measured on a multimode plate reader (CLARIOstar).

### NK cell metabolic analysis

PBMCs (1 × 10^6^ /ml) were activated using IL-15 (50ng/mL) and IL-12 (30ng/ml) or IL-18 (50ng/ml) and IL-12 (30ng/mL) for 18 hours. Cells were then labelled for extracellular markers, then fixed and permeabilized using the True-Nuclear Transcription Factor Buffer set (BioLegend) according to the manufacturer’s instructions before intracellular staining with monoclonal antibodies specific for hexokinase II (abcam) and pS6 and CD98 (Miltenyi Biotec).

### Statistics

Statistical analysis was completed using Graph Pad Prism 6 Software (USA). Data is expressed as SEM. Distribution was assessed using Sharpio-Wilk test. We determined differences between two groups using Student T-test (paired or unpaired) or Mann Whitney U test where appropriate. Analysis across 3 or more groups was performed using ANOVA with multiple measures. Correlations were determined using linear regression models and expressed using Pearson or Spearman’s rank correlation coefficient, as appropriate. P values were expressed with significance set at <0.05.

## Contributors Statement

CDB and ER performed the experiments and carried out analysis, wrote and approved the final manuscript as submitted. MS, OR, EL, HH, CMcC and DOS recruited the clinical cohorts, carried out clinical analysis and approved the final manuscript as submitted. AEH, DOS, PNM, NJ & LVS conceptualized and designed the study, analyzed the data, drafted the manuscript, and approved the final manuscript as submitted.

## Data Availability

All data is available from the corresponding author in line with ethical approval.

## Figure Legends

**Table S1:**
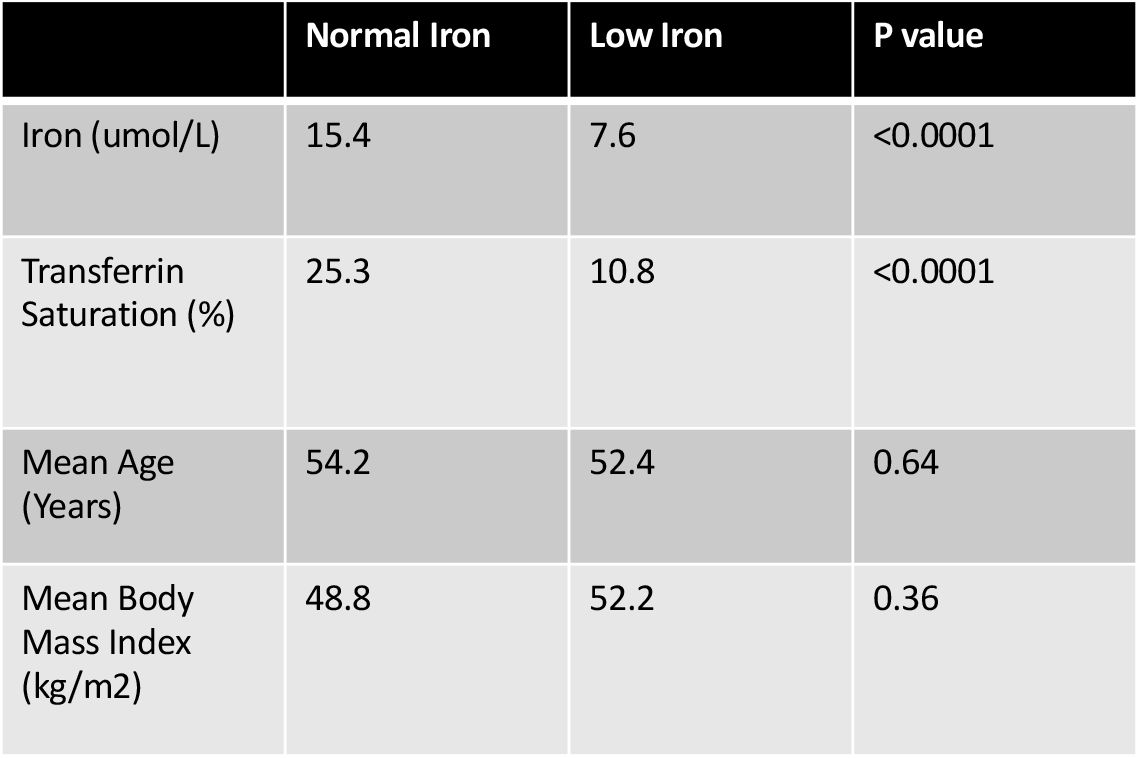
Obesity cohort demographics.

